# GENOME REPORT: Chromosome-scale genome assembly of the African spiny mouse (*Acomys cahirinus*)

**DOI:** 10.1101/2023.04.03.535372

**Authors:** Elizabeth Dong Nguyen, Vahid Nikoonejad Fard, Bernard Y. Kim, Sarah Collins, Miranda Galey, Branden R. Nelson, Paul Wakenight, Simone M. Gable, Aaron McKenna, Theo K. Bammler, Jim MacDonald, Daryl M. Okamura, Jay Shendure, David R. Beier, Jan Marino Ramirez, Mark W. Majesky, Kathleen J. Millen, Marc Tollis, Danny E. Miller

**Author notes:** These authors contributed equally (Marc Tollis, Danny E. Miller).

## Abstract

There is increasing interest in the African spiny mouse (*Acomys cahirinus*) as a model organism because of its ability for regeneration of tissue after injury in skin, muscle, and internal organs such as the kidneys. A high-quality reference genome is needed to better understand these regenerative properties at the molecular level. Here, we present an improved reference genome for *A. cahirinus* generated from long Nanopore sequencing reads. We confirm the quality of our annotations using RNA sequencing data from four different tissues. Our genome is of higher contiguity and quality than previously reported genomes from this species and will facilitate ongoing efforts to better understand the regenerative properties of this organism.

## INTRODUCTION

African spiny mice (genus *Acomys*) are a rodent species native to Africa and the Middle East. Their origin dates back to the late Miocene period ∼8.7Mya in the savannas of East Africa (Aghová *et al*. 2019). Unique adaptations to their environment have made them distinct from other rodents, as they are the first rodent species to exhibit menstruation (Bellofiore *et al*. 2017, 2021) and have the unique ability to concentrate urine to survive their arid environments (Dickinson *et al*. 2007). The African spiny mouse inhabits what is known as Evolution Canyon in lower Nahal Oren, Mount Carmel, Israel which consists of two distinct microenvironments, the hot and dry African slope and the temperate, humid and forest European slope (Hadid *et al*. 2014). The spiny mouse has thus been an evolutionary model of sympatric speciation, with populations of animals demonstrating divergence in karyotype (Volobouev *et al*. 2007), mitochondrial DNA (Hadid *et al*. 2014) and genome methylation patterns (Wang *et al*. 2022).

More recently, *Acomys cahirinus*, a member of the African spiny mouse family, has emerged as a model organism for the study of organ regeneration. Members of this family have adapted for survival in unique ways, including the ability for scarless healing of complex tissue after injury as adults. Spiny mice can shed their dorsal skin to escape the grasp of predators and then fully regenerate the lost skin without fibrotic scarring (Seifert *et al*. 2012). This scarless healing is accompanied by complete regeneration of skin including hair follicles, sebaceous glands, cartilage, adipose tissue, nerves, and blood vessels in the correct architecture for restoration of structure and function of skin tissue (Seifert *et al*. 2012; Brant *et al*. 2015; Matias Santos *et al*. 2016; Gawriluk *et al*. 2016; Jiang *et al*. 2019; Maden and Brant 2019; Harn *et al*. 2021; Brewer *et al*. 2021). The spiny mouse also demonstrates the ability to restore skeletal muscle after damage induced by cardiotoxin (Garry *et al*. 2016; Maden *et al*. 2018). These healing properties extend to internal organs; kidney damage induced using aggressive models of obstructive and ischemic injury is followed by complete regeneration of functional kidney tissue without scarring (Okamura *et al*. 2021). The spiny mouse has also been shown to exhibit resistance to myocardial ischemia and minimal scarring, as well as improvement in cardiac function after injury (Qi *et al*. 2021; Koopmans *et al*. 2021; Peng *et al*. 2021). Regeneration to this degree has been demonstrated in other mammalian species (albeit rarely), including humans, particularly in fetal tissues (Colwell *et al*. 2003; Drenckhahn *et al*. 2008; Porrello *et al*. 2011; Pratsinis *et al*. 2019; Abrams *et al*. 2021). This suggests that the potential pathways directing regeneration exist in the mammalian genome in a repressed state. A deeper understanding of the spiny mouse genome would help uncover its wound healing properties and possible reversal in nonregenerative mammalian species.

Here, we report a long-read-based chromosome-level assembly for the African spiny mouse *A. cahirinus*, a member of the *Acomys* family that is known to be capable of organ regeneration (Brewer *et al*. 2021; Okamura *et al*. 2021). The assembled *A. cahirinus* genome is more contiguous and has more complete genes when compared to previously published reference genomes (Wang *et al*. 2022), with an N50 of 127 Mb and BUSCO score of 98.5%. The *Acomys* genomic resources provided here will contribute to the better understanding of their unique organismal adaptations broadly, while accelerating further discovery of mechanisms underlying their novel adult regenerative capabilities.

## MATERIALS & METHODS

### Karyotype and banding

Chromosome analysis was performed on fibroblasts grown from ear tissue, anticoagulated blood, and bone marrow from the femur of a male *A. cahirinus*. Fibroblasts were grown to 70%– 80% confluency in DMEM/F12 with 10% FBS and 1% Pen-Strep, with rounded cells indicating active mitosis from Passage 1, 2 or 3. Anticoagulated blood was grown in RPMI (Gibco #11875093) supplemented with 10% FBS and 1% Pen-Strep and 200 µL of PHA (Gibco #10576015) for 3 days. Femurs were cut open and rinsed multiple times with 1–2 mL of RPMI supplemented with 10% FBS and 1% pen-strep. Bone marrow cells were then placed into 10- mL cultures for 24 hrs.

Samples were placed in 50 µL of Ethidium Bromide (1 mg/mL) and 50 µL of Karyomax Colcemid (10µg/mL) (Gibco #15212-012) for 1 hr. Cells were then spun down at 500g for 10 minutes. Cells were gently resuspended in 0.56% KCl and incubated at RT for 20 min. Cells spun down again at 500g for 10 min. Cells were gently resuspended in Carnoys Fixitive (3:1 Methanol:Acetic Acid) and incubated for 45 min. This was repeated twice, with incubation shortened to 10 min. Cells were then resuspended in a small volume of fresh Carnoys and dropped onto clean slides. Slides were kept at 37° for a minimum of 24 hr before banding.

For GTG banding, slides were dipped in Trypsin 2.5% (Gibco #15090-046) with NaCl for 15–60 sec, then rinsed in NaCl with FBS, then NaCl again. Slides were then stained for 10 min in Karyomax Giemsa Stain R66 Solution (Gibco #10092-013) with 50 mL of Gurr Buffer Tablets 6.8ph (Gibco #10582-013). After rinse with ddH20, slides were dried and imaged.

### Nanopore sequencing and preassembly filtering

Genomic DNA was extracted from blood from a single male *A. cahirinus* animal using a Monarch HMW DNA Extraction Kit for Cells & Blood (T3050, New England Biolabs, Ipswich MA) following the manufacturer’s recommended protocol. DNA was quantified prior to library construction using the Qubit DNA HS Assay (ThermoFischer, Waltham MA) and DNA fragment lengths were assessed using the Agilent Femto Pulse System (Santa Clara, CA). Libraries were prepared for sequencing using the Oxford Nanopore ligation kit (SQK-LSK110) following the manufacturers’ instructions, except that DNA repair and A-tailing was performed for 30 min and the ligation was allowed to continue for 1 hr. Prepared libraries were quantified using a Qubit fluorometer and 30 fmol of the library was loaded onto a Nanopore version R.9.4.1 flow cell on the PromethION platform running MinKNOW version 21.05.20. To increase output, the flow cell was washed after approximately 24 hr of sequencing then an additional 12 fmol of library was loaded and run for an additional 48 hr. Basecalling was performed using Guppy 5.0.12 (Oxford Nanopore) using the super accuracy model (dna_r9.4.1_450bps_sup_prom.cfg). Reads of quality 6 or less were discarded and NanoPlot was used to collect read statistics **(Table S1, Figure S1)**.

### Assembly and polishing

FASTQ files for assembly were extracted from unaligned bam files using samtools (Li *et al*. 2009) then Flye version 2.9 for assembly using the --nano-hq flag (Kolmogorov *et al*. 2019). Haplotigs and overlaps in the assembly were purged using purge_dups (Guan *et al*. 2020). The assembly was then polished using Medaka version 1.4.2 (https://github.com/nanoporetech/medaka) followed by a second polishing step with pilon version 1.24 (Walker *et al*. 2014). Assembly statistics at each step were generated using Quast (Gurevich *et al*. 2013) and BUSCO version 5.2.2 using the vertebrata_odb10 database (Simão *et al*. 2015) **(Table S2)**.

### Hi-C scaffolding

The primary contigs assembled from the Nanopore data were anchored to pseudo- chromosomes using 505,210,505 read pairs of a Hi-C library isolated from another *Acomys cahirinus* individual of unknown sex, downloaded from the NCBI Short Read Archive (SRX13258644) (Wang *et al*. 2022). After aligning the Hi-C reads with the ArimaHi-C Mapping Pipeline (https://github.com/ArimaGenomics/mapping_pipeline), YaHS v1.0 (Zhou *et al*. 2023) was used with default error correction for scaffolding, and Juicebox v1.11.08 (Dudchenko *et al*. 2018) was used to generate a Hi-C contact map.

### Annotation

Progressive Cactus was used (Armstrong *et al*. 2020) to perform a whole-genome alignment of the scaffolded *Acomys cahirinus* draft assembly to the *Mus musculus* GRCm39 reference genome (RefSeq GCF_000001635.27_GRCm39). Comparative annotation of the draft genomes was then performed using the Comparative Annotation Toolkit (CAT) (Fiddes *et al*. 2018). Briefly, the *M. musculus* RefSeq annotation GFF was parsed and validated with the “parse_ncbi_gff3” and “validate_gff3” programs (respectively) from CAT. The *M. musculus* reference transcript cDNA sequences were downloaded and mapped to the *M. musculus* draft genome with minimap2 (Li 2018) and provided to CAT as long-read RNA-seq reads in the “[ISO_SEQ_BAM]” field of the configuration file. For *A. cahirinus*, bulk RNA-seq data obtained from multiple pooled organs were downloaded from NCBI SRA BioProject PRJNA342864 (Bellofiore *et al*. 2017) and mapped to the draft assembly with STAR (Dobin *et al*. 2013) then provided to CAT in the “[BAMS]” field. CpG islands were identified using the cpg_lh utility from the UCSC suite of tools (Kent *et al*. 2002).

We modeled repeats *de novo* for the *Acomys cahirinus* scaffolds with RepeatModeler v2.0 (Flynn *et al*. 2020), and used RepeatMasker v4.1.3 (Smit *et al*. 1996) to (i) classify the *de novo* repeat family consensus sequences and (ii) annotate all classified repeats in the genome assembly based on the “rodentia” repeat library from RepBase v4.0.7 (Bao *et al*. 2015).

### RNA isolation and mapping

Tissues (blood, heart, liver, testis) were collected from an adult male *Acomys cahirinus* and homogenized, and RNA isolation, library generation, and sequencing were performed as previously described (Brewer *et al*. 2021; Okamura *et al*. 2021). Briefly, total RNA was extracted in Trizol solution (Ambion), DNase treated, and purified (PureLink RNA Mini Kit, Thermo Fisher Scientific). RNA was processed with KAPA’s Stranded mRNA-Seq kit (Illumina) following the manufacturer’s protocol in duplicate for each sample. The resulting libraries were assessed for library quality using fragment length and number of cycles in real-time PCR. Passing samples were sequenced on a NextSeq 500 using a 300-cycle mid-output kit, with paired 150-base pair reads.

RNA was mapped to the final assembly using bwa (version 0.7.17-r1188) (Li and Durbin 2009). Reads mapping to genomic features defined in the GTF file were counted using featureCounts using the simplified file format (Liao *et al*. 2014). RPKM was calculated for each gene using only mapped reads **(File S2)**. Venn diagrams were created in R and show overlap in genes from each tissue with RPKM values greater than 1 (**Figure 2B**).

### Comparative Genomics

#### Synteny analysis

To understand evolutionary change between the *Acomys* and *Mus* genomes, we used SynMap2 (Haug-Baltzell *et al*. 2017) on the CoGe platform (Lyons and Freeling 2008) to visualize whole-genome synteny between *Acomys cahirinus* scaffolds and the *Mus musculus* reference genome (mm39). We used lastz (Harris 2007) to map *Acomys* coding sequences to both genomes, DAGChainer (Haas *et al*. 2004) to compute chains of syntenic genes, and CodeML (Yang and Nielsen 2002) to calculate the rate of nonsynonymous (Kn) and synonymous (Ks) substitutions, as well as their ratios (Kn/Ks), with default parameters.

#### Repeat analysis

To estimate the amount of evolutionary divergence within repeat families, we generated repeat family-specific alignments using the -a flag in RepeatMasker, and calculated the average Kimura 2-parameter (K2P) sequence divergence between each annotated repeat insertion and its family consensus sequence. To correct for higher mutation rates at CpG sites, we weighted two transition mutations as 1% of a single transition. These steps were undertaken using the calcDivergenceFromAlign.pl tool in RepeatMasker. We compared the resulting repeat landscape obtained for *Acomys cahirinus* to a parallel analysis we conducted for *Mus musculus* (mm10).

#### Whole genome alignment

To further examine genomic differences between *Acomys cahirinus* and *Mus musculus*, we generated pairwise genome alignments. We first aligned *Acomys* scaffolds as queries to the mouse reference genome (mm39) with lastz (Harris 2007) using parameters K = 2400, L = 3000, Y = 9400, H = 2000, which are sensitive enough to detect orthologous exons in placental mammals (Sharma and Hiller 2017), and a default scoring matrix, followed by chaining and netting (Kent *et al*. 2003). To analyze protein-coding genes, we downloaded mm39 RefSeq gene annotations for each mouse chromosome in whole gene BED format from the UCSC Genome Browser (Kent *et al*. 2002) and used the “stitch gene blocks” tool available on Galaxy (usegalaxy.org, last accessed February 2023) to reconstruct sequence alignments for each mouse protein-coding gene ID containing the prefix “NM_” (Blankenberg *et al*. 2011). We then removed gaps in the reference alignments, removed codons with missing nucleotides which produce unknown amino acids, removed premature stop codons, and converted all filtered FASTA alignments into axt format with AlignmentProcessor.py (https://tinyurl.com/23y38664, last accessed February 2023). All alignments are included in Supplementary Data.

#### Functional analysis

To examine protein-coding differences that may point to selection pressures acting on genes since the divergence of *A. cahirinus* and *M. musculus*, we estimated the pairwise synonymous Ka and nonsynonymous Ks substitution rate, as well as the rate ratio Ka/Ks for all filtered axt gene alignments with KaKs_calculator2.0 (Wang *et al*. 2010), accounting for variable mutation rates across sites with a maximum likelihood method MS. We concatenated the results of the Ka/Ks test for each gene ID and applied the false discovery rate (FDR = 0.05) to reduce false positives (Supplementary Data). We functionally annotated all unique gene IDs with Ka/Ks > 1.0 and an adjusted p-value < 0.05 using DAVID (Sherman *et al*. 2022) and Gene Ontology enrichment (Gene Ontology Consortium 2015), applying Benjamini-Hochberg and FDR corrections to adjust for multiple testing.

### Data availability statement

RNA sequencing data and original Nanopore data are available at NCBI under bioproject PRJNA935753. The scaffolded genome assembly and gff3 files are available at https://doi.org/10.5281/zenodo.7761277. Additional annotation, alignment, and results from Ka/Ks analysis are available at https://doi.org/10.5281/zenodo.7734822.

## RESULTS & DISCUSSION

*Acomys* from our colony have a chromosomal count of 38 (19 pairs). Most autosomes are metacentric or submetacentric with a large acrocentric X, small acrocentric Y, and two pairs of small acrocentric autosomes (**Figure 1A**). These results match the *Acomys* karyotype from Morehset, Israel which is distinct from the *Acomys* karyotype generated from animals in Sinai, Egypt which have 36 chromosomes (Volobouev *et al*. 2007). Using a single male individual from our colony we generated 87.5 Gb of Nanopore data for primary assembly with a read length N50 of 63 kb and a mean read quality of 13. The initial primary assembly after purging duplicates and polishing contained 181 contigs with a contig N50 of 58.8 Mb, a longest contig of 126.8 Mb, and a total length of 2.3 Gb (**Table 1**). The contigs were anchored to 19 pseudo- chromosomes based on the Hi-C scaffolding, matching expectations from the karyotype (**Figure 1B**). Hi-C scaffolding reduced the number of assembled sequences to 129, with a scaffold N50 of 127 Mb and a total length of 2,289,268,912 bp and 79 gaps. Fifty percent of the scaffolded assembly resides on 8 scaffolds (L50). According to the BUSCO analysis of the scaffolded assembly, 98.5% of complete and partial single-copy mammalian orthologs are present, indicating a higher level of completeness than previously published reference genomes for *A. cahirinus* (Wang *et al*. 2022)

**Figure 1.**
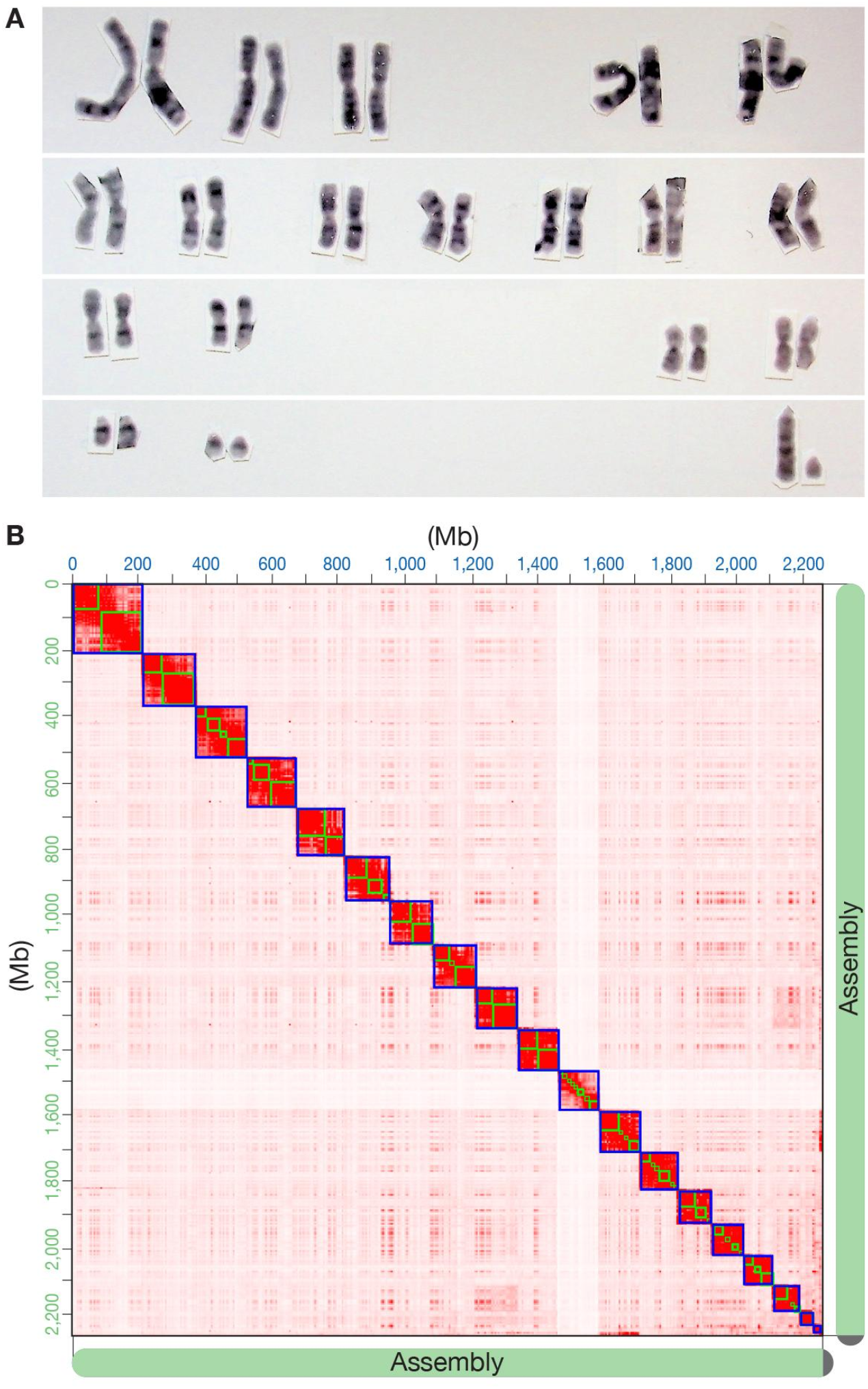
Karyotype of *Acomys cahirinus* and results of Hi-C scaffolding of the assembly. (A) Representative karyotype of a male *A. cahirinus* from the same colony that DNA and RNA were obtained from for sequencing and assembly. (**B)** Hi-C interaction contact heatmap of 19 *Acomys* pseudo-chromosomes matches the karyotype (bin size is 1 Mb).

**Figure 2.**
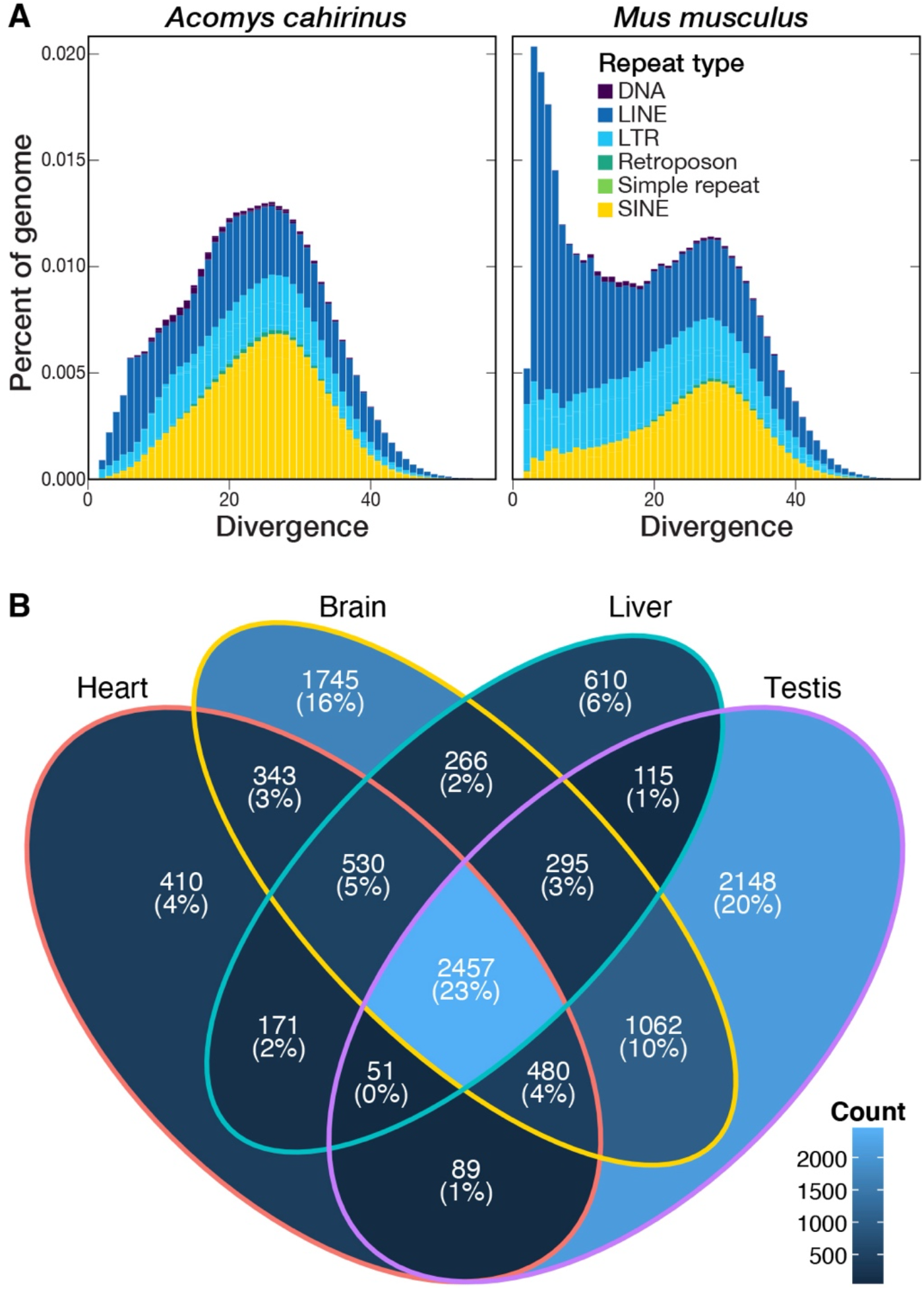
(**A**) Repeat landscapes for *Acomys cahirinus* and *Mus musculus*, visualizing the percent of each genome comprised of different types of repeats according to their divergence from family consensus (DNA, DNA transposon; LINE, long interspersed nuclear element; LTR, long terminal repeat retrotransposon; SINE, short interspersed nuclear element). (**B**) Venn diagram showing the number of genes per tissue with an RPKM value greater than 1. The majority of genes (2,457; 23%) are expressed at this level in all four tissues, while testis had the highest number of genes (2,148; 20%) with an RPKM greater than 1 not seen in other tissues.

**Table 1.**
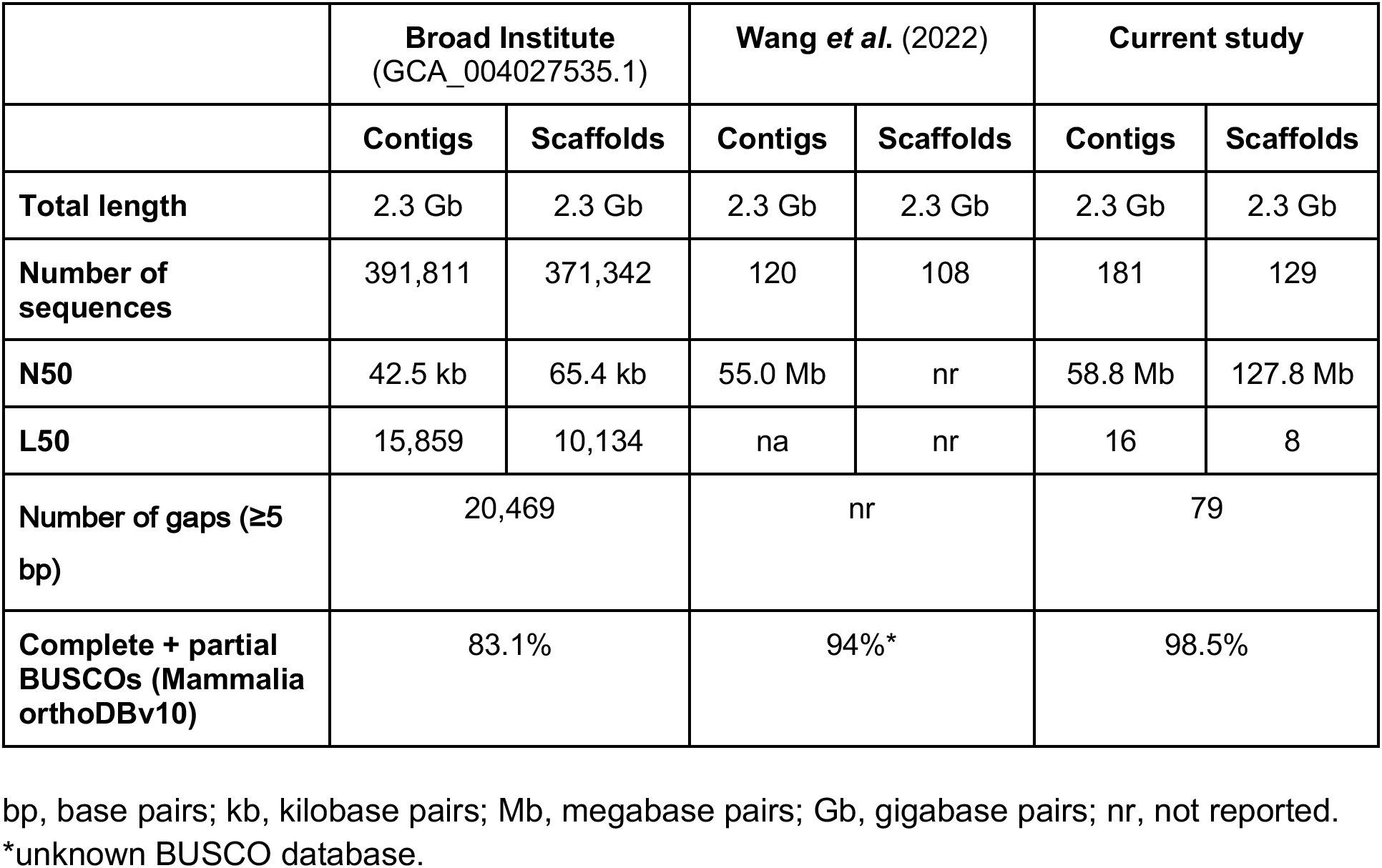
Comparison of metrics from available genome assemblies for Acomys cahirinus.

|  | **Broad Institute**<br>(GCA_004027535.1) |  | **Wang *et al.* (2022)** |  | **Current study** |  |
| --- | --- | --- | --- | --- | --- | --- |
|  | **Contigs** | **Scaffolds** | **Contigs** | **Scaffolds** | **Contigs** | **Scaffolds** |
| **Total length** | 2.3 Gb | 2.3 Gb | 2.3 Gb | 2.3 Gb | 2.3 Gb | 2.3 Gb |
| **Number of sequences** | 391,811 | 371,342 | 120 | 108 | 181 | 129 |
| **N50** | 42.5 kb | 65.4 kb | 55.0 Mb | nr | 58.8 Mb | 127.8 Mb |
| **L50** | 15,859 | 10,134 | na | nr | 16 | 8 |
| **Number of gaps (≥5 bp)** | 20,469 |  | nr |  | 79 |  |
| **Complete + partial BUSCOs (Mammalia orthoDBv10)** | 83.1% |  | 94%* |  | 98.5% |  |

bp, base pairs; kb, kilobase pairs; Mb, megabase pairs; Gb, gigabase pairs; nr, not reported.
\*unknown BUSCO database.

We estimate that approximately 37% of the *A. cahirinus* genome consists of repetitive sequences, an identical proportion to what we found in *M. musculus* (**Table 2**). Thirty four percent of the *A. cahirinus* genome consisted of interspersed repeats such as transposable elements, with most of these belonging to retrotransposons, which accounted for 30% of the genome alone. Compared to *Mus*, *Acomys* contains more SINE retrotransposons (8.3% of the genome versus 11.4%, respectively), while *Mus* contains more LINE elements (19% of the genome versus 11%, respectively). These differences can be attributed to a recent burst of LINE-1 retrotransposon activity in *M. musculus* (Sookdeo *et al*. 2013), as demonstrated by a relatively taller peak of LINE elements in the *Mus* genome at ≤10% K2P divergence compared to *Acomys* (**Figure 2A**).

**Table 2.** Repeat content of analyzed genome assemblies for Acomys cahirinus (current study) and Mus musculus (mm10)

| Parameters | Current study (*Acomys cahirinus*) | *Mus musculus* (mm10) |
| --- | --- | --- |
| Total length | 2.3 Gb | 2.7 Gb |
| GC content | 42.8% | 41.7% |
| Bases masked | 36.7% | 36.7% |
| Total interspersed repeats | 33.9% | 39% |
| Retroelements | 30.2% | 37.1% |
| SINEs | 11.7% | 8.3% |
| LINEs | 11.4% | 18.9% |
| LTR elements | 7.2% | 9.9% |
| DNA transposons | 0.63% | 0.45% |
| Unclassified | 3.0% | 1.5% |

SINEs: short interspersed nuclear elements; LINEs: long interspersed nuclear elements; LTR: long terminal repeat

Almost all forms of structural variations such as inversions, duplications, and insertions/deletions were detected in the synteny analysis between *Acomys* and *Mus* (**Figure S2**). While this suggests dynamic structural changes to the rodent genome, the mutation rate analysis indicated a major peak at Ks<<0.0, suggesting the majority of found gene pairs predate the *Mus-Acomys* divergence (**Figure S3**).

We estimated Ka/Ks for 33,197 mouse gene IDs that passed our alignment filtering. The vast majority of the genes had Ka/Ks values between 0 and 1 (mean 0.16, **Figure S4)**, indicating purifying selection acting on protein-coding genes across rodents of the Muridae family. Of the rest, 38 significant gene IDs had both Ka/Ks > 1.0 and an adjusted p-value < 0.05; 34 of these contained DAVID IDs. Three genes (*Dmrtc1b*, *Dmrtc1c1*, *Dmrtc1c2*) were annotated by Uni-Prot and InterPro as being involved in the DMRT and DMRT-like families, and 4 were enriched with the GO term meiotic cell cycle (GO:0051321). Thirty-one of the significant genes with Ka/Ks > 1.0 were enriched with PANTHER GO-Slim terms for biological processes, including spermatid development (GO:0007286, 34.2-fold enrichment, FDR = 0.0184) and germ cell development (GO:0007281, 46.9-fold enrichment, FDR = 8.59E-05). These results suggest that important differences at the amino acid level between *Mus* and *Acomys* contribute to post-speciation differences in reproductive development, which is a common result in comparative genomics analyses across mammalian species (Chai *et al*. 2021) and may be indicative of the rapid generation times and reproductive output in murid rodents.

To confirm the quality of our assembly and annotation as well as identify a broad range of expressed transcripts, we performed RNA sequencing of several tissues. We aligned short-read RNA isolated from heart, liver, brain, and testis to the assembled genome. RPKM for each annotated gene was calculated and used to determine expression levels among the four tissues for the 19,818 annotated genes. More genes with an RPKM level > 1 were observed in the brain (7,167) and testis (6,683) compared to liver (4,478) and heart (4,515) and testis had the largest number of uniquely expressed transcripts (2,148) (**Figure 2B**). This result is consistent with other studies that have demonstrated a high number of unique transcripts in brain in both *Mus musculus* and *Rattus norvegicus* (Söllner *et al*. 2017) as well as a unique number of expressed transcripts in testis (Djureinovic *et al*. 2014; Uhlén *et al*. 2015). These results provide support for the high-quality of our genome assembly and demonstrate tissue-specific expression analysis is feasible in order to better understand the regenerative capabilities of this species.

Diverse scientific disciplines have long studied *Acomys cahirinus* for their unique organismal and behavioral adaptations. Most recently, *Acomys* have emerged as an exciting and experimentally tractable adult regenerative mammalian model, as their naturally selected capacity for anti-fibrotic scarless epidermal wound healing extends across multiple internal systems and different injury contexts. Hence, our highly contiguous, high-quality genome presented here will broadly benefit the growing *Acomys* community and will accelerate more detailed investigations into the genetic and epigenetic mechanisms underlying *Acomys*’ novel capacity to maintain organ regeneration as adult mammals.

## Author Contributions

Conception: EDN, BRN, MWM, KJM, MT, DEM

Analysis: EDN, VN, SMG, BYK, AM, TKB, JM, MT, DEM

Experiments: PW, MG, AM, SC, DMO, DEM

Funding: BRN, KJM, JS, MWM, MT, DEM, DRB, JMR

Writing: EDN, BRN, KJM, MT, DEM

## Acknowledgements

We thank Angela Miller for help with editing and figure preparation.

## Funder Information

Keck Foundation for BioMedical Research (MWM, KJM)

NIH R01DK114149 (MWM)

NIH R21OD023838 (KJM, BRN)

NIH R21OD030107 (KJM)

Impetus Longevity Award (BRN)

NIH DP5OD033357 (DEM)

Brotman Baty Institute (EDN, DEM)

NIH U54CA217376 (MT)

NIH 5P50HD103524-03 (JM, TKB)

Howard Hughes Medical Institute (JS)

Seattle Research Institute Center for Developmental Biology and Regenerative Medicine (DRB)

Seattle Research Institute Center for Integrative Brain Research (JMR)

## SUPPLEMENTAL FIGURES & TABLES

**Figure S1.**
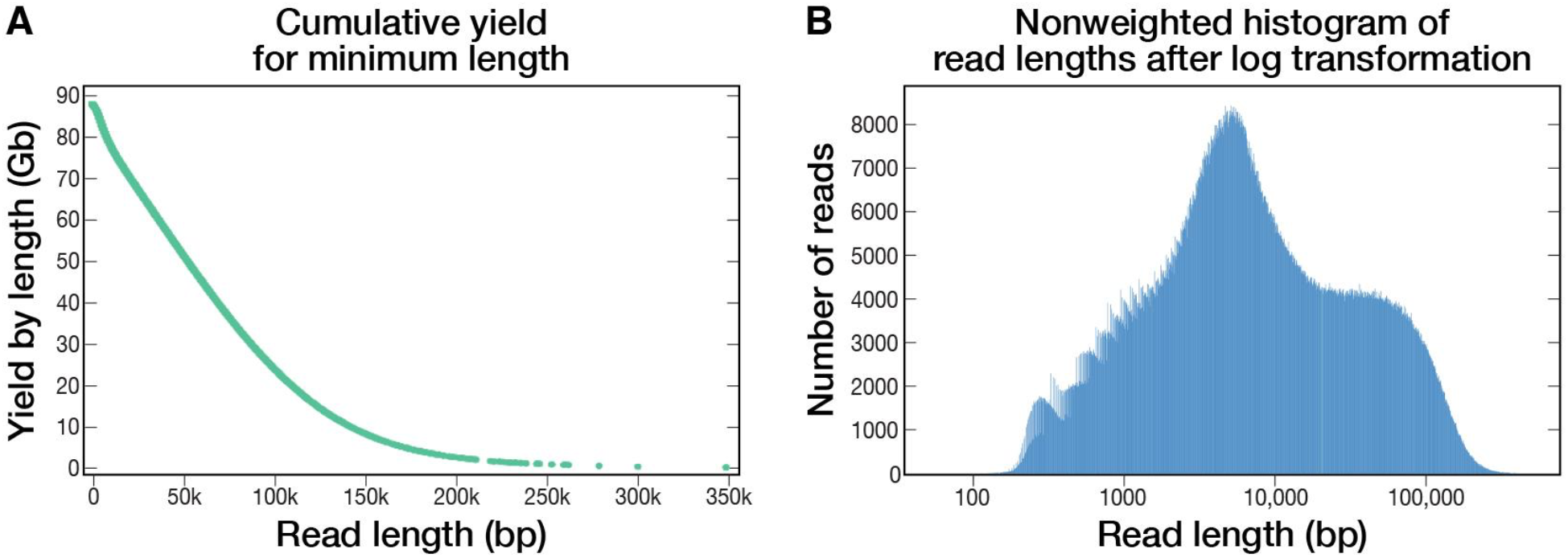
Read statistics after basecalling, generated with NanoPlot (De Coster *et al*. 2018). **(A)** Cumulative yield (in Gb) by read length. **(B)** Count of reads by length.

**Figure S2.**
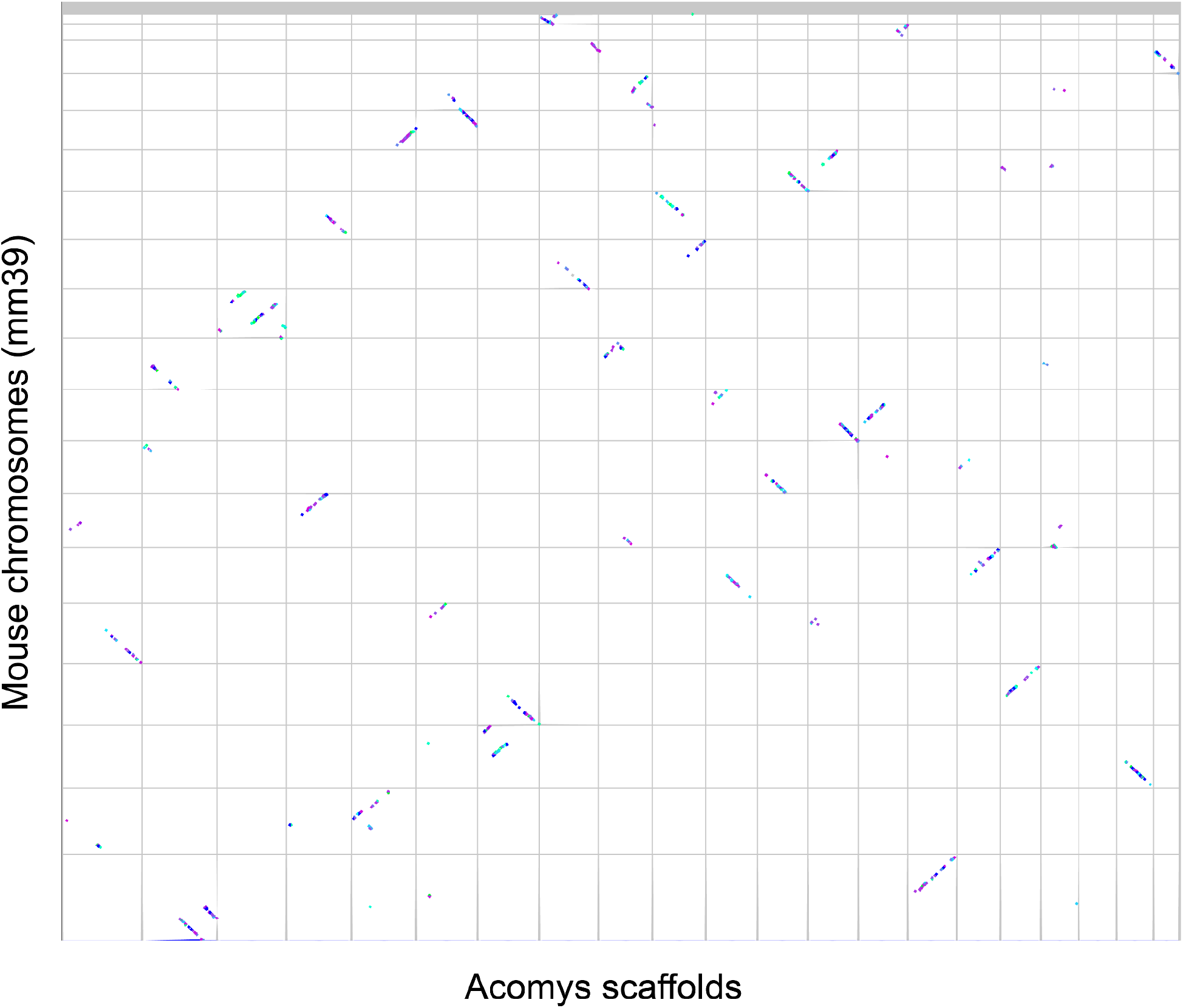
Syntenic dot plot with Kn/Ks coloration. The latest *Mus musculus* reference (mm39) is represented on the y-axis, and *Acomys cahirinus* scaffolds are on the x-axis. Grey lines separate chromosomes/scaffolds in each genome. Syntenic gene pairs are colored by the ratio of the rate of nonsynonymous to synonymous mutations (Kn/Ks), with the smallest values dark blue, followed in increasing value by violet, light blue, green, red, and orange (orange denotes ∼Kn/Ks > 1.1, which is rare in the *Acomys-Mus* comparison - see Results).

**Figure S3.**
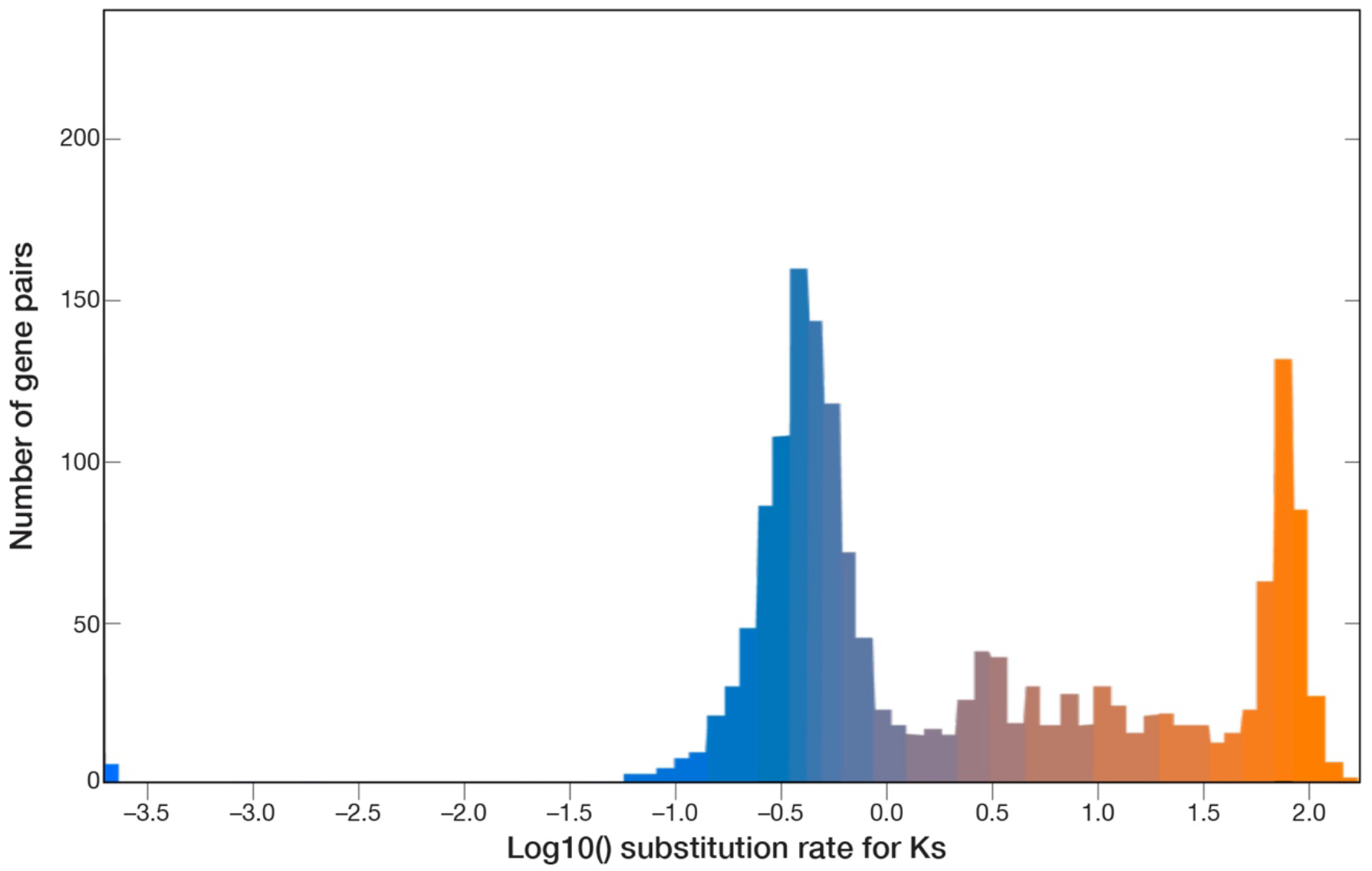
Histogram of the log10-transformed synonymous mutation (Ks) values of the syntenic gene pairs found between *Mus* (mm39) and *Acomys cahirinus*.

**Figure S4.**
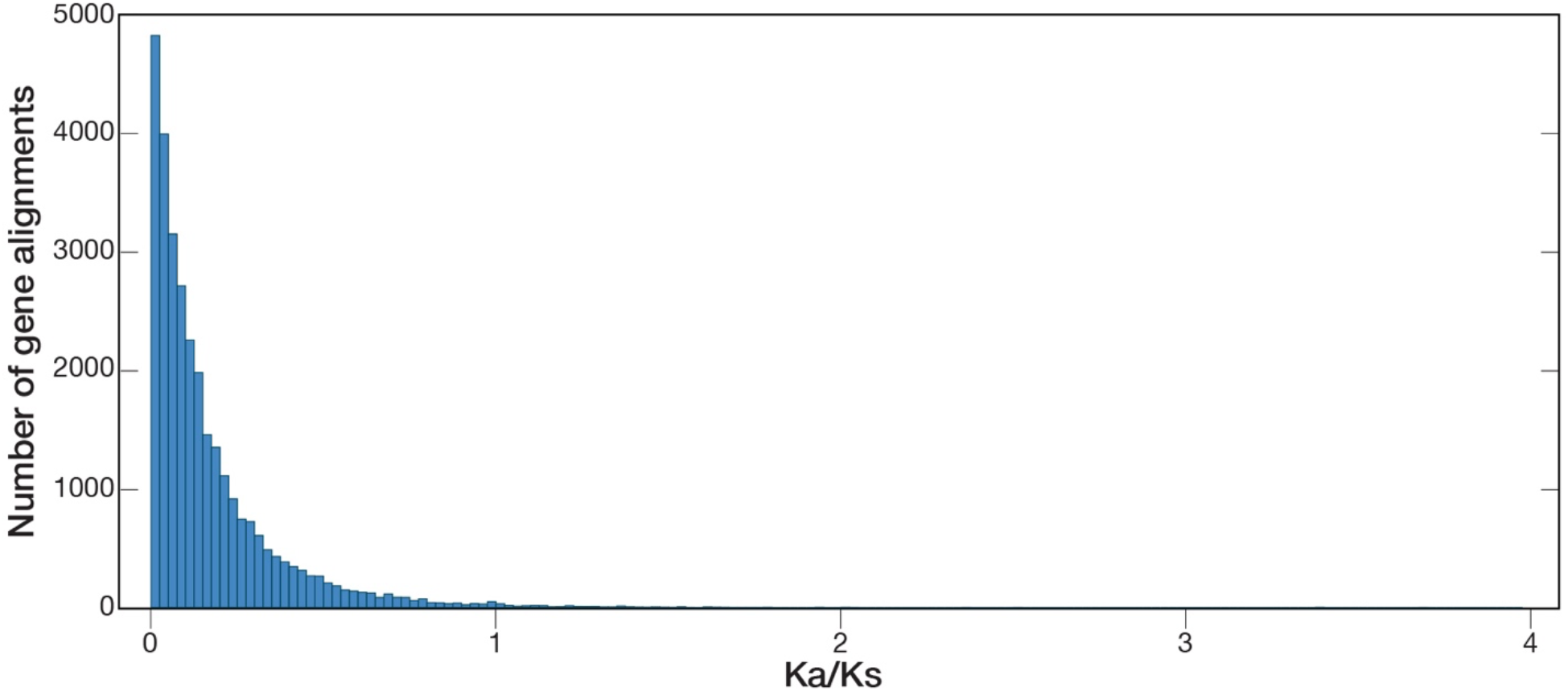
Distribution of Ka/Ks estimates for 33,197 protein-coding orthologs between *Acomys cahirinus* and *Mus musculus*. Multiple isoforms were tested for some genes.

**Table S1.** Summary of Oxford Nanopore sequencing statistics generated by NanoPlot (De Coster *et al*. 2018).

|  |  |  |
| --- | --- | --- |
| **Mean read length** | 20,158 |  |
| **Mean read quality** | 12.6 |  |
| **Median read length** | 6,129 |  |
| **Median read quality** | 12.6 |  |
| **Number of reads** | 4,345,343 |  |
| **Read length N50** | 62,867 |  |
| **STDEV read length** | 32,666 |  |
| **Total bases** | 87,593,195,281 |  |
| **Quality cutoff** | **Number of reads (%)** | **Read length** |
| >Q5 | 4,345,343 (100.0%) | 87,593.2 Mb |
| >Q7 | 4,345,166 (100.0%) | 87,592.9 Mb |
| >Q10 | 3,444,710 (79.3%) | 74,862.6 Mb |
| >Q12 | 2,483,116 (57.1%) | 59,723.1 Mb |
| >Q15 | 930,916 (21.4%) | 23,692.5 Mb |
| **Top 5 longest reads** | **Mean base call quality score** |  |
| 769,429 | 7.4 |  |
| 730,505 | 7.3 |  |
| 665,850 | 10.8 |  |
| 661,387 | 11.2 |  |
| 656,035 | 7.8 |  |

**Table S2.** Quast and BUSCO scores after each step of assembly, polishing, or scaffolding.

|  | After purging<br>duplicates | After polishing<br>with medaka | After polishing<br>with pilon | After scaffolding<br>and annotation |
| --- | --- | --- | --- | --- |
| **Quast statistics** |  |  |  |  |
| # contigs ($\geq 0$ bp) | 181 | 181 | 181 | 129 |
| # contigs ($\geq 1000$ bp) | 177 | 177 | 177 | 125 |
| # contigs ($\geq 5000$ bp) | 162 | 162 | 162 | 110 |
| # contigs ($\geq 10000$ bp) | 156 | 156 | 156 | 104 |
| # contigs ($\geq 25000$ bp) | 147 | 147 | 147 | 94 |
| # contigs ($\geq 50000$ bp) | 141 | 141 | 141 | 87 |
| Total length ($\geq 0$ bp) | 2,288,116,004 | 2,289,791,246 | 2,289,253,112 | 2,289,268,912 |
| Total length ($\geq 1000$ bp) | 2,288,113,194 | 2,289,788,406 | 2,289,250,289 | 2,289,266,089 |
| Total length ($\geq 5000$ bp) | 2,288,074,385 | 2,289,749,375 | 2,289,211,425 | 2,289,227,225 |
| Total length ($\geq 10000$ bp) | 2,288,033,368 | 2,289,707,944 | 2,289,170,056 | 2,289,185,856 |
| Total length ($\geq 25000$ bp) | 2,287,894,346 | 2,289,568,906 | 2,289,031,072 | 2,289,027,872 |
| Total length ($\geq 50000$ bp) | 2,287,707,297 | 2,289,381,919 | 2,288,844,248 | 2,288,790,854 |
| # contigs | 181 | 181 | 181 | 129 |
| Largest contig | 126,722,130 | 126,814,207 | 126,792,893 | 212,232,915 |
| Total length | 2,288,116,004 | 2,289,791,246 | 2,289,253,112 | 2,289,268,912 |
| GC (%) | 43 | 43 | 43 | 43 |
| N50 | 58,763,166 | 58,809,824 | 58,791,706 | 127,770,522 |
| N75 | 34,405,976 | 34,434,390 | 34,427,997 | 117,233,451 |
| L50 | 16 | 16 | 16 | 8 |
| L75 | 28 | 28 | 28 | 13 |
| # N's per 100 kbp | 0 | 0 | 0 | 1 |
| **BUSCO statistics** |  |  |  |  |
| Complete BUSCOs (C) | 3,290 (98.0%) | 3,300 (98.4%) | 3,305 (98.6%) | 3,304 (98.5%) |
| Complete and single-copy BUSCOs (S) | 3,218 (95.9%) | 3,225 (96.2%) | 3,232 (96.4%) | 3,233 (96.4%) |
| Complete and duplicated BUSCOs (D) | 72 (2.1%) | 75 (2.2%) | 73 (2.2%) | 71 (2.1%) |
| Fragmented BUSCOs (F) | 25 (0.7%) | 19 (0.6%) | 15 (0.4%) | 16 (0.5%) |
| Missing BUSCOs (M) | 39 (1.3%) | 35 (1.0%) | 34 (1.0%) | 34 (1.0%) |

**File S1.** Concatenated output from KaKs_calculator2.0 analysis of mouse RefSeq transcripts.

**File S2.** RPKM values for RNA sequencing data from heart, liver, brain, and testis.

## REFERENCES

Abrams, M. J., F. H. Tan, Y. Li, T. Basinger, M. L. Heithe et al., 2021 A conserved strategy for inducing appendage regeneration in moon jellyfish, Drosophila, and mice. Elife 10.:

Aghová, T., K. Palupčíková, R. Šumbera, D. Frynta, L. A. Lavrenchenko et al., 2019 Multiple radiations of spiny mice (Rodentia: Acomys) in dry open habitats of Afro-Arabia: evidence from a multi-locus phylogeny. BMC Evol. Biol. 19: 69.

Armstrong, J., G. Hickey, M. Diekhans, I. T. Fiddes, A. M. Novak et al., 2020 Progressive Cactus is a multiple-genome aligner for the thousand-genome era. Nature 587: 246–251.

Bao, W., K. K. Kojima, and O. Kohany, 2015 Repbase Update, a database of repetitive elements in eukaryotic genomes. Mob. DNA 6: 11.

Bellofiore, N., S. J. Ellery, J. Mamrot, D. W. Walker, P. Temple-Smith et al., 2017 First evidence of a menstruating rodent: the spiny mouse (Acomys cahirinus). Am. J. Obstet. Gynecol. 216: 40.e1–40.e11.

Bellofiore, N., J. McKenna, S. Ellery, and P. Temple-Smith, 2021 The Spiny Mouse-A Menstruating Rodent to Build a Bridge From Bench to Bedside. Front Reprod Health 3: 784578.

Blankenberg, D., J. Taylor, A. Nekrutenko, and Galaxy Team, 2011 Making whole genome multiple alignments usable for biologists. Bioinformatics 27: 2426–2428.

Brant, J. O., M.-C. Lopez, H. V. Baker, W. B. Barbazuk, and M. Maden, 2015 A Comparative Analysis of Gene Expression Profiles during Skin Regeneration in Mus and Acomys. PLoS One 10: e0142931.

Brewer, C. M., B. R. Nelson, P. Wakenight, S. J. Collins, D. M. Okamura et al., 2021 Adaptations in Hippo-Yap signaling and myofibroblast fate underlie scar-free ear appendage wound healing in spiny mice. Dev. Cell 56: 2722–2740.e6.

Chai, S., X. Huang, T. Wu, S. Xu, W. Ren et al., 2021 Comparative genomics reveals molecular mechanisms underlying health and reproduction in cryptorchid mammals. BMC Genomics 22: 763.

Colwell, A. S., M. T. Longaker, and H. P. Lorenz, 2003 Fetal wound healing. Front. Biosci. 8: s1240–8.

De Coster, W., S. D’Hert, D. T. Schultz, M. Cruts, and C. Van Broeckhoven, 2018 NanoPack: visualizing and processing long-read sequencing data. Bioinformatics 34: 2666–2669.

Dickinson, H., K. Moritz, E. M. Wintour, D. W. Walker, and M. M. Kett, 2007 A comparative study of renal function in the desert-adapted spiny mouse and the laboratory-adapted C57BL/6 mouse: response to dietary salt load. Am. J. Physiol. Renal Physiol. 293: F1093–8.

Djureinovic, D., L. Fagerberg, B. Hallström, A. Danielsson, C. Lindskog et al., 2014 The human testis-specific proteome defined by transcriptomics and antibody-based profiling. Mol. Hum. Reprod. 20: 476–488.

Dobin, A., C. A. Davis, F. Schlesinger, J. Drenkow, C. Zaleski et al., 2013 STAR: ultrafast universal RNA-seq aligner. Bioinformatics 29: 15–21.

Drenckhahn, J.-D., Q. P. Schwarz, S. Gray, A. Laskowski, H. Kiriazis et al., 2008 Compensatory growth of healthy cardiac cells in the presence of diseased cells restores tissue homeostasis during heart development. Dev. Cell 15: 521–533.

Dudchenko, O., M. S. Shamim, S. S. Batra, N. C. Durand, N. T. Musial et al., 2018 The Juicebox Assembly Tools module facilitates de novo assembly of mammalian genomes with chromosome-length scaffolds for under $1000. bioRxiv 254797.

Fiddes, I. T., J. Armstrong, M. Diekhans, S. Nachtweide, Z. N. Kronenberg et al., 2018 Comparative Annotation Toolkit (CAT)-simultaneous clade and personal genome annotation. Genome Res. 28: 1029–1038.

Flynn, J. M., R. Hubley, C. Goubert, J. Rosen, A. G. Clark et al., 2020 RepeatModeler2 for automated genomic discovery of transposable element families. Proc. Natl. Acad. Sci. U. S. A. 117: 9451–9457.

Garry, G. A., M. L. Antony, and D. J. Garry, 2016 Cardiotoxin Induced Injury and Skeletal Muscle Regeneration. Methods Mol. Biol. 1460: 61–71.

Gawriluk, T. R., J. Simkin, K. L. Thompson, S. K. Biswas, Z. Clare-Salzler et al., 2016 Comparative analysis of ear-hole closure identifies epimorphic regeneration as a discrete trait in mammals. Nat. Commun. 7: 11164.

Gene Ontology Consortium, 2015 Gene Ontology Consortium: going forward. Nucleic Acids Res. 43: D1049–56.

Guan, D., S. A. McCarthy, J. Wood, K. Howe, Y. Wang et al., 2020 Identifying and removing haplotypic duplication in primary genome assemblies. Bioinformatics 36: 2896–2898.

Gurevich, A., V. Saveliev, N. Vyahhi, and G. Tesler, 2013 QUAST: quality assessment tool for genome assemblies. Bioinformatics 29: 1072–1075.

Haas, B. J., A. L. Delcher, J. R. Wortman, and S. L. Salzberg, 2004 DAGchainer: a tool for mining segmental genome duplications and synteny. Bioinformatics 20: 3643–3646.

Hadid, Y., T. Pavlícek, A. Beiles, R. Ianovici, S. Raz et al., 2014 Sympatric incipient speciation of spiny mice Acomys at “Evolution Canyon,” Israel. Proc. Natl. Acad. Sci. U. S. A. 111: 1043–1048.

Harn, H. I.-C., S.-P. Wang, Y.-C. Lai, B. Van Handel, Y.-C. Liang et al., 2021 Symmetry breaking of tissue mechanics in wound induced hair follicle regeneration of laboratory and spiny mice. Nat. Commun. 12: 2595.

Harris, R. S., 2007 Improved pairwise alignment of genomic DNA.

Haug-Baltzell, A., S. A. Stephens, S. Davey, C. E. Scheidegger, and E. Lyons, 2017 SynMap2 and SynMap3D: web-based whole-genome synteny browsers. Bioinformatics 33: 2197– 2198.

Jiang, T.-X., H. I.-C. Harn, K.-L. Ou, M. Lei, and C.-M. Chuong, 2019 Comparative regenerative biology of spiny (Acomys cahirinus) and laboratory (Mus musculus) mouse skin. Exp. Dermatol. 28: 442–449.

Kent, W. J., R. Baertsch, A. Hinrichs, W. Miller, and D. Haussler, 2003 Evolution’s cauldron: duplication, deletion, and rearrangement in the mouse and human genomes. Proc. Natl. Acad. Sci. U. S. A. 100: 11484–11489.

Kent, W. J., C. W. Sugnet, T. S. Furey, K. M. Roskin, T. H. Pringle et al., 2002 The human genome browser at UCSC. Genome Res. 12: 996–1006.

Kolmogorov, M., J. Yuan, Y. Lin, and P. A. Pevzner, 2019 Assembly of long, error-prone reads using repeat graphs. Nat. Biotechnol. 37: 540–546.

Koopmans, T., H. van Beijnum, E. F. Roovers, A. Tomasso, D. Malhotra et al., 2021 Ischemic tolerance and cardiac repair in the spiny mouse (Acomys). NPJ Regen Med 6: 78.

Li, H., 2018 Minimap2: pairwise alignment for nucleotide sequences. Bioinformatics 34: 3094– 3100.

Liao, Y., G. K. Smyth, and W. Shi, 2014 featureCounts: an efficient general purpose program for assigning sequence reads to genomic features. Bioinformatics 30: 923–930.

Li, H., and R. Durbin, 2009 Fast and accurate short read alignment with Burrows–Wheeler transform. Bioinformatics 25: 1754–1760.

Li, H., B. Handsaker, A. Wysoker, T. Fennell, J. Ruan et al., 2009 The Sequence Alignment/Map format and SAMtools. Bioinformatics 25: 2078–2079.

Lyons, E., and M. Freeling, 2008 How to usefully compare homologous plant genes and chromosomes as DNA sequences. Plant J. 53: 661–673.

Maden, M., and J. O. Brant, 2019 Insights into the regeneration of skin from Acomys, the spiny mouse. Exp. Dermatol. 28: 436–441.

Maden, M., J. O. Brant, A. Rubiano, A. G. W. Sandoval, C. Simmons et al., 2018 Perfect chronic skeletal muscle regeneration in adult spiny mice, Acomys cahirinus. Sci. Rep. 8: 8920.

Matias Santos, D., A. M. Rita, I. Casanellas, A. Brito Ova, I. M. Araújo et al., 2016 Ear wound regeneration in the African spiny mouse Acomys cahirinus. Regeneration (Oxf) 3: 52–61.

Okamura, D. M., C. M. Brewer, P. Wakenight, N. Bahrami, K. Bernardi et al., 2021 Spiny mice activate unique transcriptional programs after severe kidney injury regenerating organ function without fibrosis. iScience 24: 103269.

Peng, H., K. Shindo, R. R. Donahue, E. Gao, B. M. Ahern et al., 2021 Adult spiny mice (Acomys) exhibit endogenous cardiac recovery in response to myocardial infarction. NPJ Regen Med 6: 74.

Porrello, E. R., A. I. Mahmoud, E. Simpson, J. A. Hill, J. A. Richardson et al., 2011 Transient regenerative potential of the neonatal mouse heart. Science 331: 1078–1080.

Pratsinis, H., E. Mavrogonatou, and D. Kletsas, 2019 Scarless wound healing: From development to senescence. Adv. Drug Deliv. Rev. 146: 325–343.

Qi, Y., O. Dasa, M. Maden, R. Vohra, A. Batra et al., 2021 Functional heart recovery in an adult mammal, the spiny mouse. Int. J. Cardiol. 338: 196–203.

Seifert, A. W., S. G. Kiama, M. G. Seifert, J. R. Goheen, T. M. Palmer et al., 2012 Skin shedding and tissue regeneration in African spiny mice (Acomys). Nature 489: 561–565.

Sharma, V., and M. Hiller, 2017 Increased alignment sensitivity improves the usage of genome alignments for comparative gene annotation. Nucleic Acids Res. 45: 8369–8377.

Sherman, B. T., M. Hao, J. Qiu, X. Jiao, M. W. Baseler et al., 2022 DAVID: a web server for functional enrichment analysis and functional annotation of gene lists (2021 update). Nucleic Acids Res. 50: W216–21.

Simão, F. A., R. M. Waterhouse, P. Ioannidis, E. V. Kriventseva, and E. M. Zdobnov, 2015 BUSCO: assessing genome assembly and annotation completeness with single-copy orthologs. Bioinformatics 31: 3210–3212.

Smit, A. F. A., R. Hubley, and P. Green, 1996 Repeat-Masker Open-3.0. 1996–2010 <http://www.repeatmasker.org>.

Söllner, J. F., G. Leparc, T. Hildebrandt, H. Klein, L. Thomas et al., 2017 An RNA-Seq atlas of gene expression in mouse and rat normal tissues. Sci Data 4: 170185.

Sookdeo, A., C. M. Hepp, M. A. McClure, and S. Boissinot, 2013 Revisiting the evolution of mouse LINE-1 in the genomic era. Mob. DNA 4: 3.

Uhlén, M., L. Fagerberg, B. M. Hallström, C. Lindskog, P. Oksvold et al., 2015 Proteomics. Tissue-based map of the human proteome. Science 347: 1260419.

Volobouev, V., J. C. Auffray, V. Debat, C. Denys, J. C. Gautun et al., 2007 Species delimitation in the Acomys cahirinus–dimidiatus complex (Rodentia, Muridae) inferred from chromosomal and morphological analyses. Biol. J. Linn. Soc. Lond. 91: 203–214.

Walker, B. J., T. Abeel, T. Shea, M. Priest, A. Abouelliel et al., 2014 Pilon: an integrated tool for comprehensive microbial variant detection and genome assembly improvement. PLoS One 9: e112963.

Wang, Y., Z. Qiao, L. Mao, F. Li, X. Liang et al., 2022 Sympatric speciation of the spiny mouse from Evolution Canyon in Israel substantiated genomically and methylomically. Proc. Natl. Acad. Sci. U. S. A. 119: e2121822119.

Wang, D., Y. Zhang, Z. Zhang, J. Zhu, and J. Yu, 2010 KaKs_Calculator 2.0: a toolkit incorporating gamma-series methods and sliding window strategies. Genomics Proteomics Bioinformatics 8: 77–80.

Yang, Z., and R. Nielsen, 2002 Codon-substitution models for detecting molecular adaptation at individual sites along specific lineages. Mol. Biol. Evol. 19: 908–917.

Zhou, C., S. A. McCarthy, and R. Durbin, 2023 YaHS: yet another Hi-C scaffolding tool. Bioinformatics 39.:

